# BAP1 and YY1 regulate expression of death receptors in malignant pleural mesothelioma

**DOI:** 10.1101/2020.08.31.274951

**Authors:** Y Ishii, KK Kolluri, A Pennycuick, E Nigro, D Alrifai, E Borg, M Falzon, K Shah, N Kumar, SM Janes

## Abstract

Malignant pleural mesothelioma (MPM) is a rare, aggressive, and incurable cancer arising from the mesothelial lining of the lungs with few treatment options. We recently reported loss of function of the nuclear deubiquitinase BRCA associated protein-1 (BAP1), a frequent event in MPM, is associated with sensitivity to tumour necrosis factor-related apoptosis-inducing ligand (TRAIL). As a potential underlying mechanism, here we report that BAP1 negatively regulates the expression of TRAIL receptors: death receptors 4 (DR4) and 5 (DR5). Using tissue microarray (TMAs) of tumour samples from MPM patients, we found a strong inverse correlation between BAP1 and TRAIL receptors. *BAP1* knockdown increased DR4 and DR5 expression, whereas overexpression of BAP1 had the opposite effect. Reporter assays confirmed wild-type *BAP1*, but not catalytically-inactive mutant *BAP1*, reduced promoter activities of *DR4* and *DR5*, suggesting deubiquinase activity plays an important role in the regulation of gene expression. Co-IP studies demonstrated direct binding of BAP1 and the transcription factor Ying Yang 1 (YY1) and ChIP assays revealed BAP1 and YY1 to be enriched in the promoter regions of *DR4* and *DR5*. Notably, shRNA knockdown of *YY1* also increased DR4 and DR5 expression, and sensitivity to TRAIL. These results demonstrate that BAP1 and YY1 together negatively regulate transcriptional activity of TRAIL receptors. BAP1 and YY1 may both therefore be strong therapeutic targets to enhance the efficacy of TRAIL-induced apoptosis.

**Statement of significance:** We describe how the most-frequently mutated tumour suppressor gene in mesothelioma regulates the response to TNF-related apoptosis-inducing ligand (TRAIL). These findings will accelerate a biomarker-driven cancer therapy.

## Introduction

Malignant pleural mesothelioma (MPM) is a rare, aggressive cancer that arises from the mesothelial lining of the lungs and is commonly associated with occupational exposure to asbestos. There are currently no curative therapies. Standard first line treatment is combination chemotherapy consisting of an anti-folate and a platinum agent which offers only a modest survival benefit (1). Advances in the understanding of MPM tumour biology have led to the development of multiple novel targeted agents currently in preclinical and clinical development. Many of these therapies lack a biomarker for activity and results so far have not delivered an effective clinical therapy (2).

A molecular target of significant interest in MPM is *BRCA associated protein 1* (*BAP1)*(3–5). *BAP1* mutations are frequent in MPM (23-67%) and in other tumour types including uveal melanoma (31-50%), cholangiocarcinoma (20-25%) and clear cell renal cell carcinoma (CCRCC) (8-14%) (6–19). BAP1 is a nuclear deubiquitinase (DUB) that binds to a number of transcription factors through which it regulates gene transcription and modulates cellular pathways (4,5). The response to drugs that act upon these pathways, including PARP and EZH2 inhibitors, has been shown to be increased in the absence of BAP1 function (20). Clinical trials of these drugs in *BAP1* mutant MPM are underway (21).

Our group has previously presented data to support that loss of BAP1 function results in sensitivity to the death receptor (DR) agonist recombinant tumour necrosis factor-related apoptosis-inducing ligand (rTRAIL) (22). TRAIL is a member of the tumour necrosis factor (TNF) cytokine superfamily. It activates the extrinsic apoptotic pathway by binding to either of two death receptors, DR4 or DR5, which leads to the recruitment of the adaptor protein FADD and caspase-8 to form the death-inducing signalling complex (DISC) (23). Once formed, catalytic subunits of caspase-8 are cleaved and activate downstream effector caspases that result in apoptosis (24,25). Activation of this pathway by TRAIL is specific to cancer cells, however the mechanism of this selectivity is poorly understood (26,27). Several therapeutic DR agonists including rTRAIL and agonistic DR4/5 antibodies have been developed (28–30). Clinical trials of such agents to date have demonstrated broad tolerability, but unfortunately limited therapeutic benefit (31). Potential reasons include the suboptimal pharmacokinetics of compounds, resistant cell populations and the lack of a targeting biomarker (32). Novel DR agonists with improved pharmacokinetics are in development and potential biomarkers such as BAP1 are emerging (33,34).

Whilst we have extensively validated the association between loss of BAP1 function and increased sensitivity to rTRAIL within *in vitro, in vivo* and *ex vivo* models, the mechanism underlying this association remains unclear (22). Here we aim to delineate this mechanism. We hypothesise that BAP1 activity modulates expression of proteins of the extrinsic and intrinsic apoptosis pathways with an increase in pro-apoptotic protein expression in the absence of BAP1 activity. We identify both BAP1 activity and rTRAIL sensitivity correlate with expression of the death receptors DR4 and DR5 at the transcriptional level. As BAP1 is a transcriptional regulator without DNA binding sites, we searched for the transcriptional factor that cooperates with BAP1 to modulate expression of DR4 and DR5 identifying the polycomb group (PcG) protein YY1.

## Results

### Loss of BAP1 activity correlates with increased DR4 and DR5 expression and increased rTRAIL sensitivity

We have previously shown that MPM cells with loss of BAP1 function are more sensitive to treatment with rTRAIL (22). To determine the mechanism underlying this, we investigated the expression of death receptors DR4 and DR5, whose levels have known to significantly contribute to the sensitivity (35,36), and nuclear BAP1 expression, a surrogate for *BAP1* wild-type status (7). By immunohistochemical analysis using human tissue micro arrays (TMAs) (88 cores from 32 patients) (Fig. 1A), we observed a significant correlation between loss of nuclear BAP1 expression to higher DR4 and DR5 expression (Fig. 1B and Supplementary Fig. S1). This correlation was further supported by immunohistochemistry in primary mesothelioma tissue samples collected as part of the MSO1 clinical trial (NCT00075699) (37). MPM tissue samples that lacked nuclear BAP1 did show elevated levels of DR4 and DR5 (Supplementary Fig. S2B and S2C). Interestingly, using cytokeratin 5 (CK5) and calretinin to confirm MPM, we observed that DR4 and DR5 are expressed in MPM cells but not surrounding stromal tissue (Supplementary Fig. S2A). Such differential expression of death receptors is an existing theory for the selectivity of rTRAIL and other death receptor agonists for cancer cells (35).

**Figure 1:**
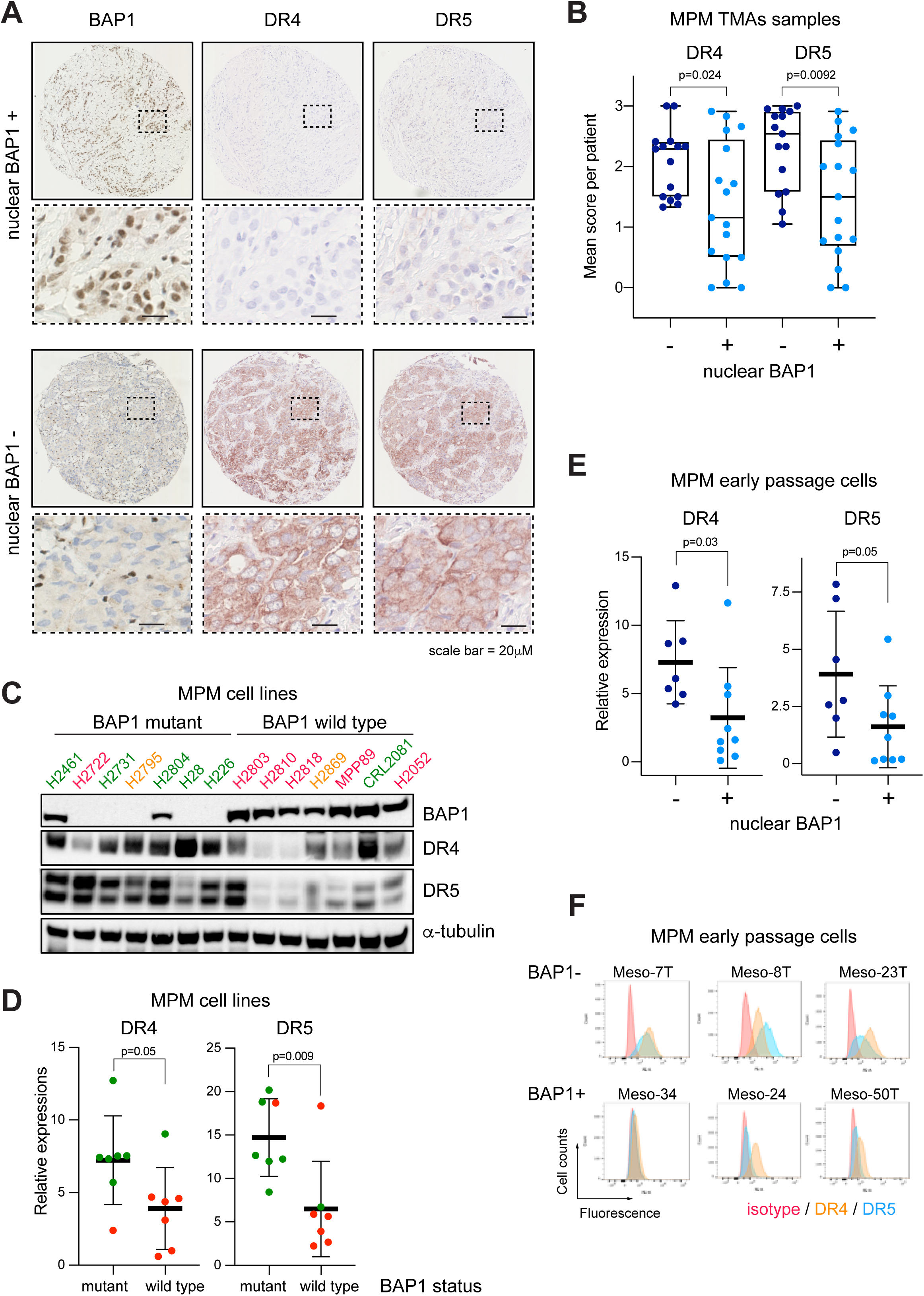
Expression levels of DR4 and DR5 are inversely correlated with BAP1 expression in malignant pleural mesothelioma. **A**, Representative images of IHC of DR4 and DR5 in a core from an MPM tissue microarray (TMA) with and without nuclear BAP1 expression (from 88 cores of 32 patients). **B**, Semi-quantitative analysis of DR4 and DR5 expression in MPM TMA cores with (n=42) and without (n=46) nuclear BAP1 expression. Each dot represents an average score per patient. t-test; p=0.024 (DR4) and p=0.0092 (DR5). See method section for details. **C**, Immunoblots of DR4, DR5 and BAP1 protein expression in BAP1 mutant (n=7) vs BAP1 wild-type (n=7) MPM cell lines. Duplet bands of DR5 represent two splice variants, DR5-short (DR5-S) and DR5-long (DR5-L). Sensitivity to rTRAIL treatment is indicated as font colour: green sensitive (S); orange partially sensitive (PS); red resistant (R). **D**, Quantitative analysis of immunoblot intensity of DR4 and DR5 in wild type BAP1 and mutant BAP1 MPM cell lines (DR4 t-test; p=0.05, DR5 t-test; p=0.008). Sensitivity to rTRAIL treatment is indicated as font colour as indicated in (C). **E**, Quantitative analysis of immunoblot intensity of DR4 and DR5 in early-passage MPM cells with (+) and without (-) nuclear BAP1 expression. (DR4 t-test; p=0.03, DR5 t-test; p=0.05). **F**, Flow cytometry analysis of DR4 and DR5 cell surface expression in early passage MPM cells with (BAP1+) and without (BAP1-) nuclear BAP1 expression alongside an isotype control.

Immunoblot analysis of a panel of MPM cell lines (7 *BAP1* mutant, 7 *BAP1* wild-type) also showed a strong inverse correlation between BAP1 and DR4 and DR5 expression (Fig. 1C). Quantitative analysis of immunoblots confirmed an association between the mutational status of *BAP1* in MPM cells and DR4 and DR5 expression; *BAP1* mutant cell lines expressed significantly higher levels of both DR4 and DR5 (Fig. 1D). We observed additional heterogeneous changes in expression across 20 other proteins involved in the extrinsic and intrinsic apoptosis pathways, however, they were all unrelated to the mutational status of *BAP1* or rTRAIL sensitivity (Supplementary Fig. S3). In addition to comparing steady-state levels of anti-apoptotic proteins, we also compared their expression after treatment with rTRAIL, as TRAIL has been documented in some cells to induce anti-apoptotic, rather than pro-apoptotic, pathways (38–42). We examined c-FLIP, a catalytically inactive caspase-8 homologue that competes with caspase-8, inhibitors of apoptosis proteins (cIAP1/2), MAPK and NFκB pathways that enhance proliferation and induce cIAPs (35). We saw no induction of these proteins, excluding this as a mechanism of TRAIL-resistance in BAP1-mutant cells (Supplementary Fig. S4). Furthermore, we conducted immunoblot analysis of human early passage, unsequenced MPM cultures (MesobanK UK) (43). We have previously shown that strong nuclear BAP1 expression is highly correlated with rTRAIL resistance in these cells (22). Immunoblot analysis revealed that DR4 and DR5 expression was higher in the nuclear BAP1-negative group (Supplementary Fig. S5). Quantitative analysis of immunoblots confirmed the association between nuclear BAP1 expression and DR4 and DR5 expression (Fig. 1E). Flow cytometry analysis also showed higher surface expression of DR4 and DR5 in early passage MPM cultures with loss of nuclear BAP1 expression (Fig. 1F). Taken together, our data demonstrate strong inverse correlations between BAP1 expression and DR4 and DR5 expression, which may underlie the ability of BAP1 to determine rTRAIL sensitivity.

### Loss of BAP1 function increases DR4 and DR5 expression in malignant but not in non-transformed cells

To further investigate the relationship between BAP1 and DR4 and DR5 expression we knocked down BAP1 expression in a *BAP1*-wild type MPM cell line using a lentiviral shRNA construct. *BAP1* knockdown significantly increased expression of both DR4 and DR5 (Fig. 2A). Notably, induction of cleaved caspase-8 and cleaved PARP was observed only in BAP1 knockdown cells indicating apoptosis activation only in the absence of BAP1. We confirmed BAP1 knockdown resulted in increased sensitivity to rTRAIL in these cells (Fig. 2B). To examine the effect across additional tumour types we next knocked down *BAP1* in two *BAP1* wild-type clear cell renal cell carcinoma (CCRCC) cell lines. This also resulted in increased expression of DR4 and DR5 and increased sensitivity to rTRAIL (Fig. 2C and D). BAP1 knockdown further resulted in increased DR4 and DR5 mRNA levels (Fig. 2E) indicating the effect of BAP1 on DR4 and DR5 expression is at the transcriptional level. Significantly, *BAP1* knockdown in human lung fibroblasts and human bronchial epithelial cells (HBECs) did not result in increased expression of DR4 and DR5 or increased sensitivity to rTRAIL suggesting this effect is specific to malignant cells (Fig. 2F and G).

**Figure 2:**
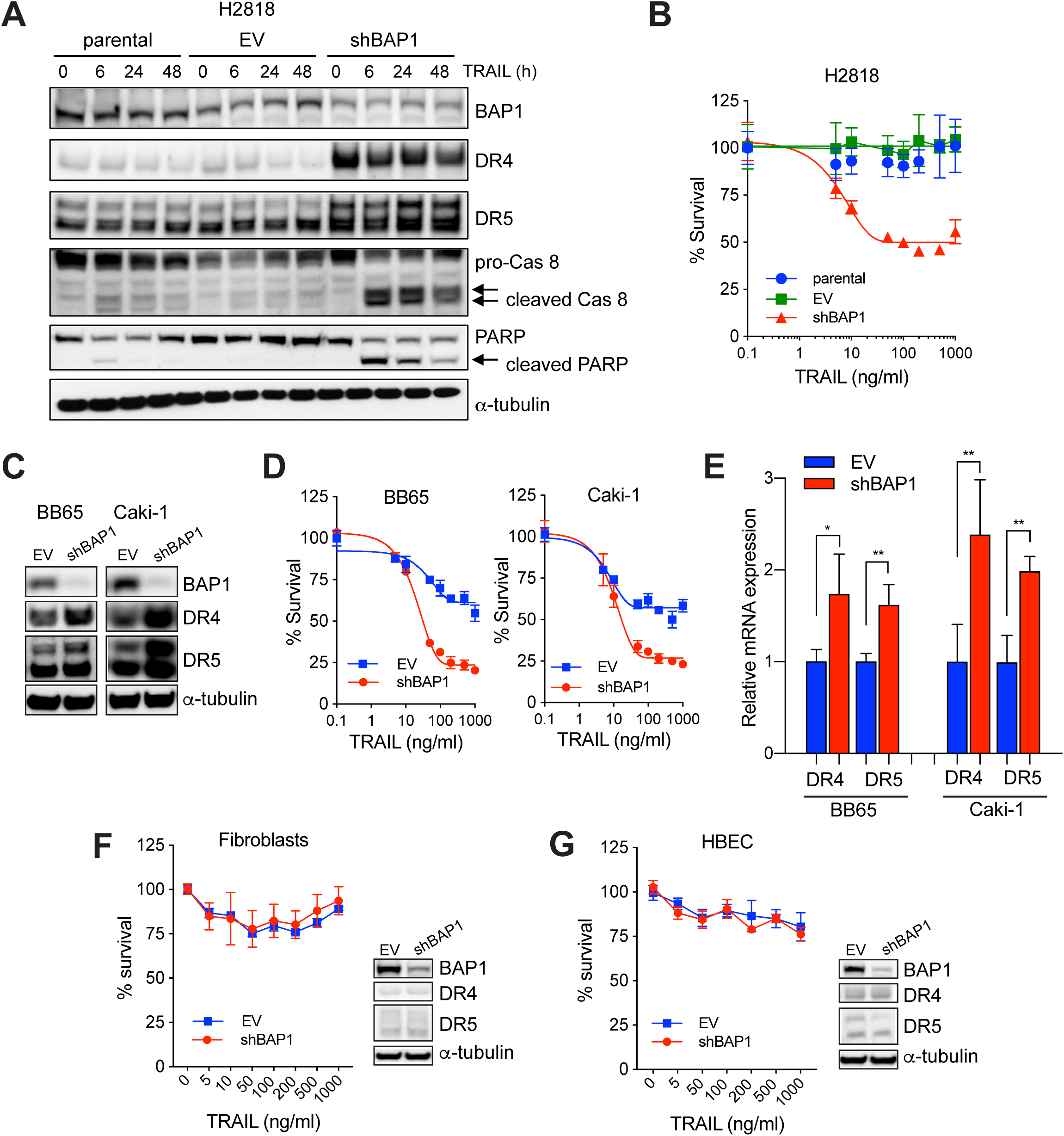
BAP1 knockdown increases death receptors expression and TRAIL sensitivity in cancer cells. **A**, Immunoblots of pro-apoptotic proteins in parental, BAP1 shRNA (shBAP1) or empty vector shRNA (EV) transduced BAP1-wild-type MPM cells (H2818) across multiple time points (0, 6, 12, 24 and 48 hours) post rTRAIL treatment (100ng/mL). **B**, Cell viability assay of parental, shBAP1- or EV-transduced H2818 cells following treatment with a dose range of rTRAIL (0-1000ng/ml) for 72 hours. **C**, Immunoblots of BAP1, DR4 and DR5 in BAP1-wild-type-clear cell renal cell carcinoma (CCRCC) cells (BB65 and Caki-1) transduced with BAP1 (shBAP1) or empty vector (EV) shRNA. **D**, Cell viability assay of EV- or shBAP1-transduced BAP1 wild-type CCRCC cells following treatment with a dose range of rTRAIL (0-1000ng/ml) for 72 hours. **E**, Relative expression of DR4 and DR5 mRNA in CCRCC cells transduced with EV or shBAP1 assessed by qPCR. Relative mRNA expression was normalized to beta-2-micro-globlin (B2M) expression. Data shown are the mean ± s.d. of two experiments performed in triplicates. t-test; *p<0.05, **p<0.01. **F & G**, Cell viability assay of fibroblasts (**F**) and human bronchial epithelial cells (HBECs) (**G**) transduced with EV or shBAP1 following treatment with a dose range of rTRAIL (0-1000ng/ml) for 72 hours. Immunoblots of BAP1, DR4 and DR5 in these cells are also shown. Error bars represent the standard deviation.

### BAP1 negatively regulates transcription of *DR4* and *DR5*

We next transduced a *BAP1*-null early passage MPM cell line, Meso-8T, with a lentiviral construct expressing wild-type *BAP1* (wt-BAP1) or *BAP1* with an inactivating mutation in the deubiquitinase site (C91A-BAP1 or A95D-BAP1). In accordance with *BAP1* knockdown data, transduction with wt-BAP1 but not C91A-BAP1 resulted in a decrease in DR4 and DR5 expression (Fig. 3A). Flow cytometry confirmed a decrease in surface expression of DR4 and DR5 in cells transduced with wt-BAP1 but not C91A-BAP1 or A95D-BAP1 (Fig. 3B). Cell survival assays confirmed transduction with wt-BAP1, but not C91A-BAP1 or A95D-BAP1, resulted in a significant reduction in rTRAIL sensitivity (Fig. 3C). Concordantly, we saw decreased activation of caspase-8, caspase-3 and reduced PARP cleavage in wt-BAP1 transduced relative to C91A-BAP1 transduced cells (Fig. 3A) reflective of reduced apoptotic pathway activation. Quantitative PCR analysis demonstrated that DR4 and DR5 mRNA expression were both decreased in cells transduced with wt-BAP1 relative to those transduced with C91A-BAP1 suggesting regulation of DR4 and DR5 expression by catalytically active BAP1 is at the transcriptional level (Fig. 3D). These results were confirmed in a further MPM cell line, H28 (Supplementary Fig. S6). Subsequently we tested the effect of BAP1 on *DR4* and *DR5* transcription more directly. Meso-8T cells were transduced with lentiviral vectors with luciferase reporters under the control of *DR4* or *DR5* promoters (44). These reporter cells were also transduced with either wt-BAP1 or A95D-BAP1. Cells transduced with wt-BAP1 displayed a significantly lower luciferase activity than those transduced with A95D-BAP1 or the parental cell line reflecting decreased *DR4* and *DR5* transcriptional activity in the presence of functional BAP1 (Fig. 3E).

**Figure 3:**
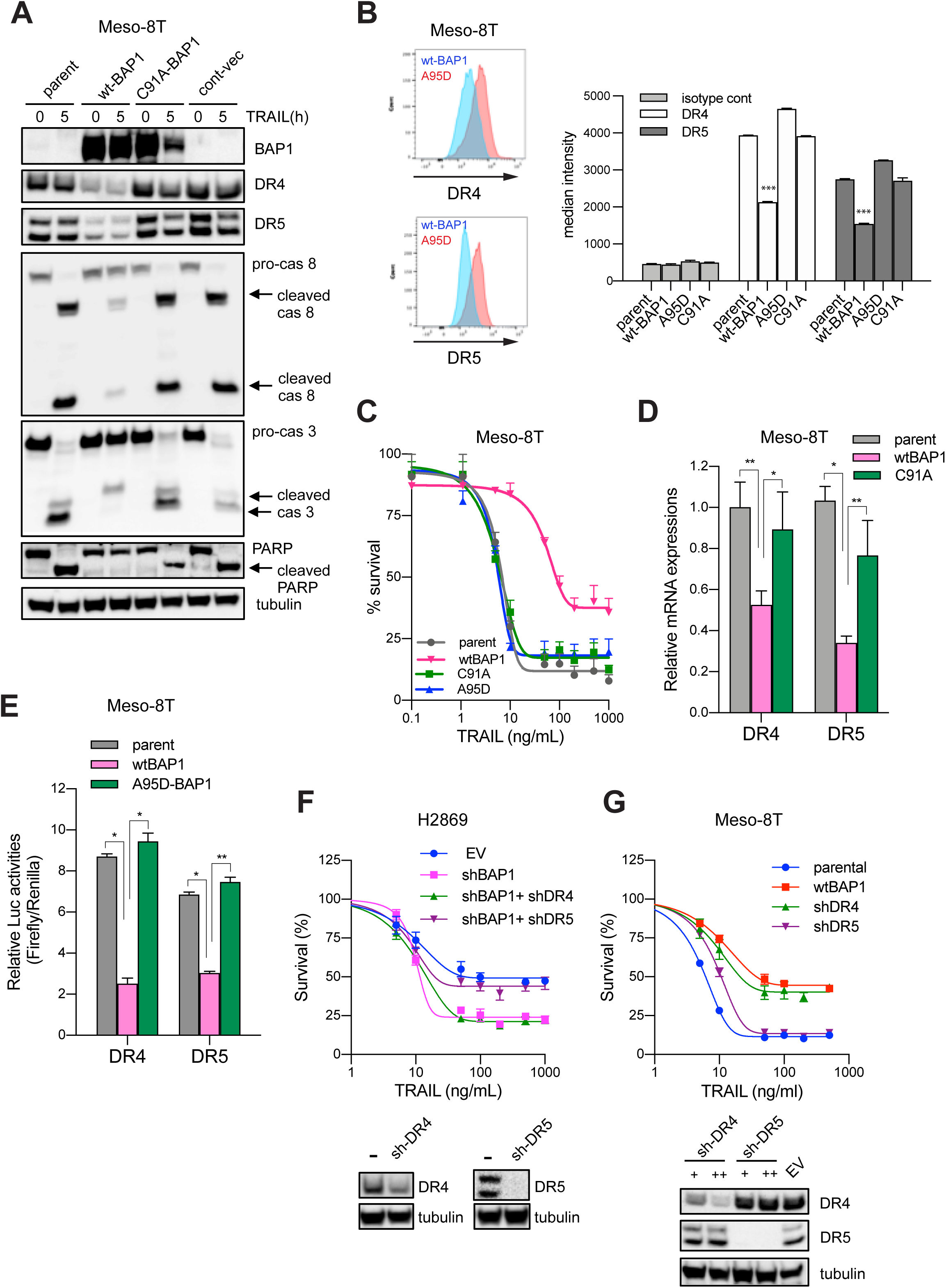
The deubiquitinase function of BAP1 regulates the transcription of DR4 and DR5. **A**, Immunoblots of pro-apoptotic proteins in BAP1 null early passage mesothelioma cells (Meso-8T) transduced with constructs expressing wild type-BAP1 (wt-BAP1), deubiquitinase mutant BAP1 (C91A-BAP1) or a control vector (cont-vec) untreated and after 5 hours of rTRAIL treatment (50 ng/mL). **B**, Flow cytometry analysis of cell surface expression of DR4 and DR5 expression in Meso-8T cells transduced with constructs expressing wild type-BAP1 (wt-BAP1) or one of two deubiquitinase mutant BAP1 vectors (C91A or A95D). One-way ANOVA; ***p<0.001. **C**, Cell viability assay of Meso-8T cells transduced with wt-BAP1 or one of two deubiquitinase mutant BAP1 vectors (C91A or A95D) following treatment with a dose range of rTRAIL (0-1000ng/ml) for 72 hours. **D**, Relative DR4 and DR5 mRNA expression in parental Meso-8T cells and cells transduced with wt-BAP1 or C91A-BAP1. Relative mRNA expression was normalized to beta-2-microgloblin (B2M) expression. Data are shown as the mean ± s.d. of two experiments performed in triplicates. *, P<0.05; **, P<0.01. **E**, Reporter assay for promoter activities of DR4 and DR5 in parental Meso-8T cells transduced with a luciferase reporter under the control of DR4 or DR5 promoter and cells further transduced with wt-BAP1 or A95D-BAP1. Firefly luciferase/Renilla luciferase ratios were determined as relative luciferase activities. Data are shown as the mean ± s.d. of two experiments (n=6 in each experiment). *, P<0.05; **, P<0.01. **F**, Cell viability assay of BAP1 wild-type H2869 cells transduced with EV or shBAP1 following treatment with a dose range of rTRAIL (0-1000ng/ml) for 72 hours and for shBAP1 cells further transduced with DR4 (shDR4) or DR5 shRNA (shDR5) following the same treatment. **G**, Cell viability assay of parental BAP1 null Meso-8T early passage MPM cells transduced with wild-type BAP1 (wtBAP1) or DR4 (shDR4) or DR5 shRNA (shDR5) following treatment with a dose range of rTRAIL (0-1000ng/ml) for 72 hours. +,++; lentiviral titer. Error bars represent the standard deviation.

Together the above results support that the deubiquitinase activity of BAP1 mediates transcriptional repression of *DR4* and *DR5*. To determine whether this in turn determines rTRAIL sensitivity, we used two complementary approaches. First, we knocked down *DR4* or *DR5* in *BAP1* wild-type H2869 MPM cells transduced with BAP1 shRNA. BAP1 knockdown increased the sensitivity of H2869 cells to rTRAIL as expected (Fig. 3F). Interestingly, DR5, but not DR4 knockdown, in shBAP1-H2869 cells abolished the effect of BAP1 knockdown, resulting in rTRAIL resistance (Fig. 3F). Second, we knocked down *DR4* or *DR5* in the BAP1-null, rTRAIL-sensitive Meso-8T cell line. *DR5* knockdown only slightly decreased rTRAIL sensitivity but *DR4* knockdown reduced it to a similar level as transduction with wild-type *BAP1* (Fig. 3G). It has been reported that some cells preferentially or exclusively use one of the two death receptors to induce apoptosis, although little is known about the mechanism (30). For example, haematological cancers seem to prefer DR4 for induction of apoptosis (45,46), whereas solid tumours appear to exhibit heterogeneity in death receptor preference (30,47,48).

### YY1 negatively regulates transcription of *DR4* and *DR5*

As BAP1 does not bind to DNA directly (5), we aimed to identify transcription factors that bind to the promoter regions of DR4 and DR5. Bioinformatic analysis of 2000 nucleotides of the promoter region of DR4 and DR5 was conducted. From candidates identified (Supplementary Fig. S7), YY1 was selected for further analysis as it has previously been shown to negatively regulate *DR5* expression in prostate cancer (49,50). Furthermore, YY1 has been shown to bind directly to BAP1 forming a polycomb repressor-deubiquitinase (PR-DUB) complex capable of regulating gene expression (51). shRNA knockdown of *YY1* in MPM and CCRCC cells resulted in increased expression of both DR4 and DR5 without affecting steady-state levels of BAP1 (Fig. 4A). shRNA knockdown of *BAP1* in MPM cells also did not affect steady-state levels of YY1. In addition, we did not observe any difference in YY1 expression based on BAP1 mutational status and BAP1 expression level (Supplementary Fig. S8). qPCR analysis confirmed increased mRNA expression of *DR4* and *DR5* in cells transduced with YY1 shRNA (Fig. 4B). *YY1* knockdown also significantly increased sensitivity to rTRAIL and the DR5 agonist Medi3039 in MPM and CCRCC cells (Fig. 4C)(52).

**Figure 4:**
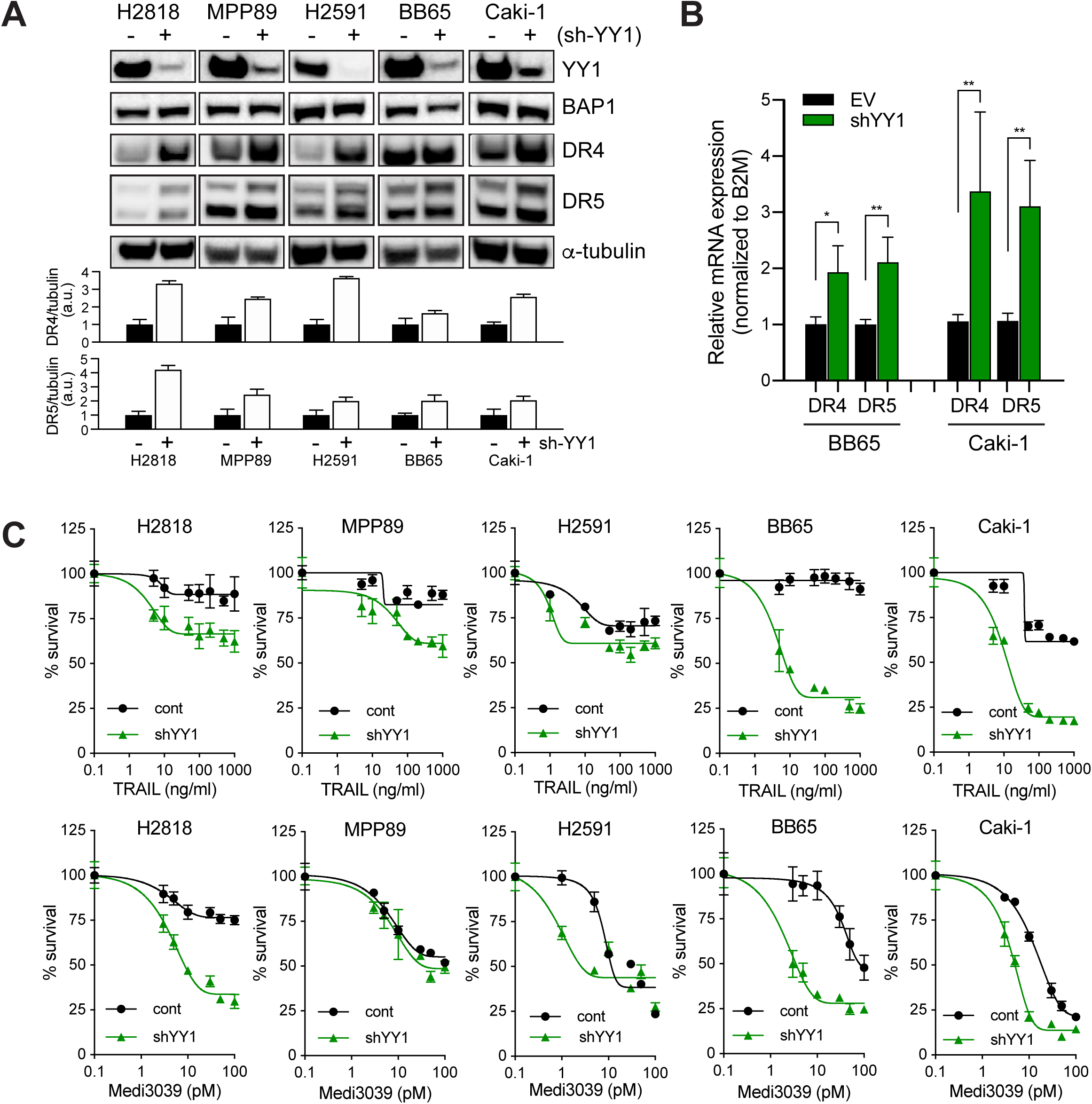
YY1 knockdown increases the expression of death receptors and rTRAIL-induced cell death. **A**, Immunoblots of YY1, BAP1, DR4 and DR5 in BAP1-wild-type MPM cells (H2818, MPP89 and H2591) or CCRCC cells (BB65 and Caki-1) transduced with YY1 shRNA (+) or an empty vector shRNA (-). Quantitative analysis of DR4 and DR5 bands from three independent experiments was performed. Average data after normalisation to tubulin were shown as bar graphs. **B**, Relative DR4 and DR5 mRNA expression in CCRCC cells (Caki-1 and BB65) transduced with YY1-shRNA or EV-shRNA. Relative mRNA expression was normalized to beta-2-microgloblin (B2M) expression. Data are shown as the mean ± s.d. of two experiments performed in triplicates. *, P<0.05; **, P<0.01. **C**, Cell viability assays of BAP1 wild-type MPM and CCRCC cells transduced with EV shRNA or shYY1 following treatment with a dose range of rTRAIL (0-1000ng/ml) or MEDI3039 (0.1-100pM) for 72 hours. Error bars represent the standard deviation.

### YY1 recruits BAP1 to DR4 and DR5 promoters to facilitate transcriptional repression

BAP1 has been shown to form a ternary complex with YY1 and HCF-1 (Host Cell Factor 1) in HeLa cells (51). Therefore, we aimed to determine if BAP1 and YY1 also directly bind in MPM and CCRCC cells. Protein extracts from H2818, MPP89 and Caki-1 cells were co-immunoprecipitated (co-IP) using anti-YY1 antibody or IgG as a control. Immunoblot confirmed the interaction of endogenous YY1 with BAP1 (Fig. 5A). To verify the specificity of these results, we compared results of co-IP assay in BAP1-null MPM cells (Meso-8T) that were transduced with wt-BAP1 or control vector alone. A strong interaction of YY1 and BAP1 was detected only in cells transduced with wt-BAP1 but not control vector, confirming the specificity of the YY1/BAP1 interaction (Fig. 5B).

**Figure 5:**
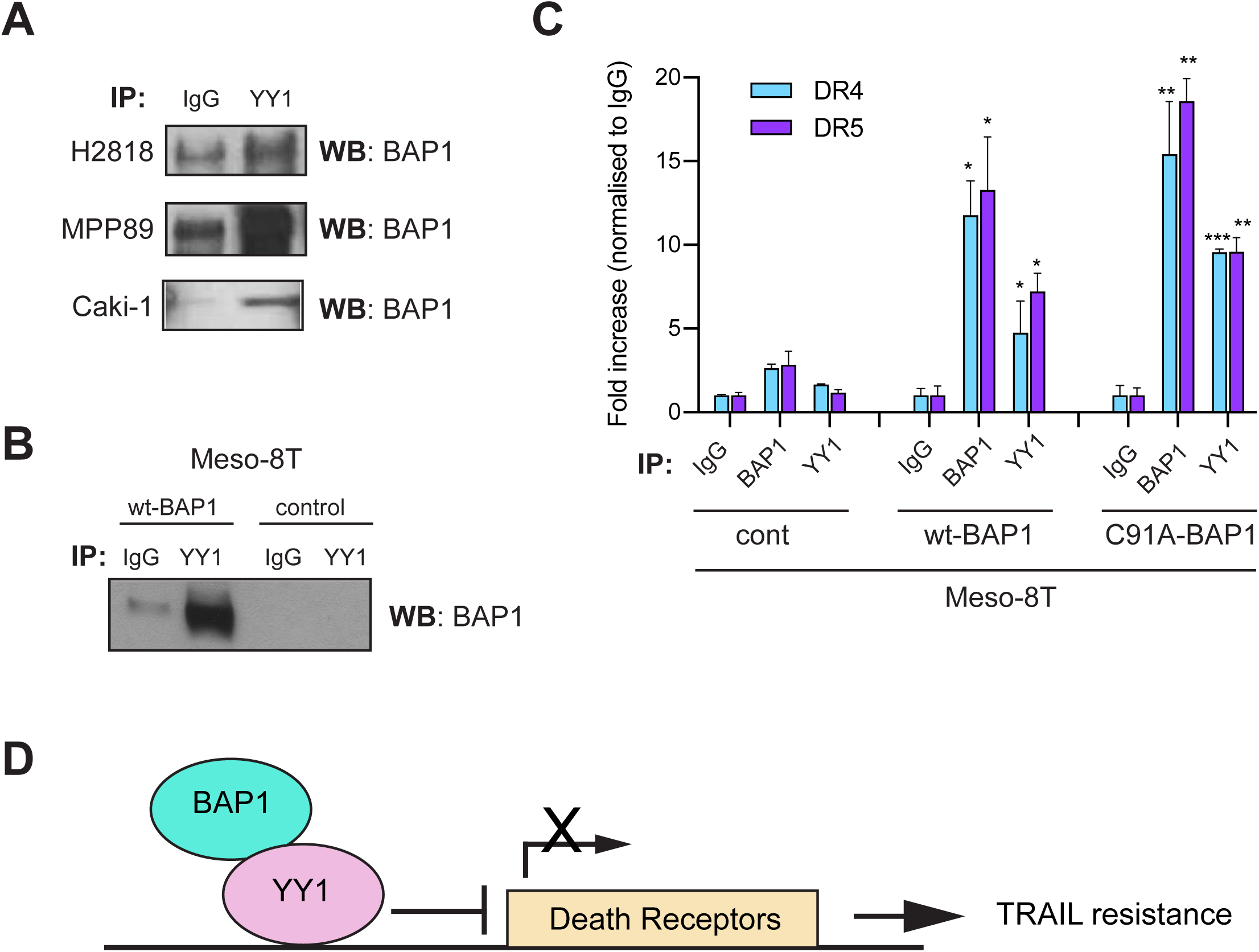
YY1 recruits BAP1 to the promoter regions of DR4 and DR5 and represses their transcriptional activities. **A**, Co-Immunoprecipitation (Co-IP) of endogenous YY1 and BAP1 in MPM (H2818, MPP89) and CCRCC (Caki-1) cells. **B**, Co-IP of YY1 and BAP1 in BAP1 null early passage MPM Meso-8T cells transduced with wild-type BAP1 (wt-BAP1) or a control vector. **C**, Enrichment of BAP1 and YY1 in the promoter regions of DR4 and DR5. Meso-8T cells were overexpressed with wt-BAP1, catalytically-inactive mutant-BAP1 (C91A-BAP1) or a control vector (cont). Chromatin Immunoprecipitation (ChIP) was performed against BAP1, YY1 or IgG control followed by qPCR using primers specific for promoter regions of DR4 or DR5. Error bars represent the standard deviation. P-values are calculated to compare against IgG control using student t-test (n=3); *p<0.05, **p<0.01. **D**, Schematic model of the transcriptional regulation of TRAIL death receptors by BAP1 and YY1.

In addition to physical interactions, we sought to examine the functional interaction between YY1 and BAP1. As BAP1 does not have a DNA binding domain, but directly interacts with the transcriptional repressor YY1, we hypothesized that YY1 recruits BAP1 to the promoter regions of DR4 and DR5. Chromatin immunoprecipitation (ChIP) assays were performed with antibodies for BAP1, YY1 or IgG as a control. The immunoprecipitated DNA was analysed with probes for DR4 or DR5 by qPCR in Meso-8T cells transduced with wt-BAP1, C91A-BAP1 or a control vector. Both BAP1 and YY1 were enriched in the promoter regions of DR4 and DR5 in cells transduced with wt-BAP1 but not the control vector (Fig. 5C). Interestingly, BAP1 and YY1 were also enriched in these promoter regions in cells transduced with C91A-BAP1 indicating YY1 recruits BAP1 regardless of deubiquitinase activity. This finding is consistent with previous reports that catalytically inactive BAP1 is also recruited to FoxK2-binding regions (53). Catalytically inactive BAP1 has also previously been shown to form a complex with YY1 (51).

## Discussion

We have recently reported that loss of BAP1 function is a predictive biomarker for rTRAIL sensitivity in cancer (22). In this study, we delineate the underlying molecular mechanism. We demonstrate BAP1 is recruited to the promoter regions of *DR4* and *DR5* by the transcriptional regulator YY1. This complex facilitates transcriptional repression of *DR4* and *DR5* and requires BAP1 deubiquitinase activity. Decreased cell surface expression of DR4 and DR5 and reduced activation of the apoptotic pathway in turn mediates rTRAIL resistance in *BAP1* wild-type cells. Conversely, increased cell surface expression of DR4 and DR5 in *BAP1* mutant cells mediates the observed increased sensitivity to rTRAIL. Various mechanisms of resistance to rTRAIL and other death receptor agonists have been suggested (54). Evidence supports that low expression of DR4 and DR5 due to mutations, promoter methylation, constitutive endocytosis or deficient transport to the cell surface is important (54–57). Indeed, strategies to enhance the efficacy of rTRAIL treatment, such as a combination with chemotherapeutic drugs, have been demonstrated to mediate these effects through increased death receptor expression (24). Our results are consistent with these data and support the centrality of death receptor expression in TRAIL therapeutics.

YY1 inhibition has previously been shown to upregulate DR5 expression and enhance rTRAIL sensitivity in prostate cancer and B-non-Hodgkin lymphoma cells (49,50,58). Here however we show that YY1 is involved in the transcriptional regulation of both DR4 and DR5 and that binding to BAP1 and BAP1 deubiquitinase activity is also required. BAP1 is known to be a powerful transcriptional regulator and it’s precise targets dictated by the protein partners with which it complexes (5). Notably, YY1 has previously been shown to recruit BAP1 and the transcriptional cofactor HCF-1 to the cox7 promoter, a component of the mitochondrial respiratory chain, where it facilitates transcriptional activation of *cox7* (51). An additional cofactor may direct BAP1 and YY1 to the DR4 and DR5 promoters. We have previously shown that mutation of the ASXL binding site on BAP1 and ASXL1 knockdown also increases rTRAIL sensitivity (22). The BAP1/ASXL1 complex is a polycomb repressor deubiquitinase complex capable of deubiquitination of histone 2A, a process which modulates expression of the polycomb genes (5). Interestingly, YY1 has also been shown to interact with polycomb proteins (51,59). It may therefore be that YY1 interacts with both BAP1 and ASXL1 to modulate death receptor expression.

YY1 and BAP1 may be involved more widely in the transcriptional regulation of the TNF receptor superfamily. Nitric oxide has been shown to inhibit YY1 binding to the *Fas* promoter resulting in Fas upregulation and cell sensitisation to Fas-ligand induced apoptosis in prostate cancer (60). YY1 has also been shown to supress the Fas promoter activity in B-non-Hodgkin lymphoma and colon cancer (61,62). We have also previously demonstrated that BAP1 knockdown sensitises MPM cells to Fas ligand, CH11 and TNF-alpha (22).

Our findings have significant clinical utility. We have already proposed BAP1 expression to be a stratifying biomarker for sensitivity to death receptor agonists (22). In addition to BAP1, the current study also demonstrates that YY1 knockdown enhances the sensitivity to TRAIL and DR5 agonist. YY1 is overexpressed in many types of cancer and high expression correlates with poor clinical outcomes and resistance to chemotherapy and immunotherapy making it an attractive therapeutic target (63,64). Thus, targeting the BAP1/YY1 axis may be an additional novel therapeutic strategy in TRAIL therapeutics.

## Materials and Methods

### Cell Culture

All cancer cell lines were obtained from the Wellcome Trust Sanger Institute except the H226 line that was kindly gifted from Dr. P. Szlosarek (Barts Cancer Institute). Cancer cell lines were cultured in RPMI-1640, Dulbecco’s modified Eagle’s medium (DMEM) or DMEM and nutrient mix 12 medium (DMEM:F12) supplemented with 10% fetal bovine serum (FBS), penicillin/ streptavidin and sodium pyruvate. Early passage human mesothelioma cells were purchased from MesobanK (43) and cultured in RPMI-1640 medium supplemented with 5% FBS, 25 mM HEPES, penicillin/ streptavidin and sodium pyruvate. Primary human lung fibroblasts (kind gift from Dr. R. Chambers at UCL) were cultured in DMEM media supplemented with 10% FBS and penicillin/streptavidin in an incubator with 10% CO_2_ (65). Experiments were conducted on cells between passage 6 and 8. Primary human bronchial epithelial cells (HBECs) were obtained from endobronchial biopsies with patient consent as previously described (66). Ethical approval was obtained through the National Research Ethic Committee (REC reference 06/Q0505/12). HBECs were cultured in bronchial epithelial growth medium (BEGM; Lonza) on top of 3T3-J2 mouse embryonic fibroblast feeder cells inactivated by mitomycin-C treatment (0.4 μg/ml, Sigma-Aldrich).

### XTT cell viability assay

Cells were seeded in 96-well plates in 100 µl media per well at a density of 40,000 cells/ml one day prior to treatment with soluble recombinant TRAIL (rTRAIL; Peprotech-310-04) or MEDI3039 (Medimmune). XTT reagent and the activation solution (Applichem, A8088) were mixed and added to the cells at the end of treatment. The plate was returned to a CO_2_ incubator to incubate for 2 hours, the absorbance at a wavelength of 490nm was measured using a microplate reader. Relative cell viability was calculated as a fraction of viable cells relative to untreated cells.

### Immunoblotting

Cells were lysed in RIPA lysis buffer (Sigma-Aldrich) with protease inhibitors (Complete-mini; Roche) on ice to extract protein. 30 µg of protein samples were separated by SDS–PAGE and transferred onto nitrocellulose membranes using iBlot2 Dry Blotting System (Thermo Fisher Scientific). Membranes were incubated with specific primary antibodies, washed, incubated with secondary antibodies and visualised using an ImageQuant LAS 4000 imaging system (GE Healthcare). A list of antibodies used for immunoblotting is provided in Supplementary Table1. Quantification of bands was performed using ImageJ (Image Processing and Analysis in Java).

### Immunoprecipitation (IP)

Cells were lysed in IP buffer containing 0.2% NP-40, 20mM Tris-HCl (pH 7.4), 150mM NaCl, 10% glycerol and protease inhibitors. The lysates were incubated overnight with gentle rocking with anti-YY1 antibody (Abcam: ab38422) or IgG (Cell Signaling: 2729). Protein-A magnetic beads (Pierce Biotechnology Inc.) were added and incubation was continued for 1 hour. The beads were washed with IP buffer and proteins eluted from the beads by heating with SDS sample buffer. Proteins were separated by SDS-PAGE and immunoblotting was performed as described above with anti-BAP1 antibody (Santa Cruz: sc-28383).

### Plasmids

Full-length *BAP1* cDNA was amplified by PCR from pCMV6-AC *BAP1* plasmid (Origene-SC117256) and cloned into the lentiviral plasmid pCCL-CMV-flT vector. Vectors expressing mutant *BAP1* constructs were generated by site-directed mutagenesis (New England Biolabs E0554) of the pCCL-CMV-BAP1 vector as previously described (22).

### RNA interference

Short hairpin RNAs (shRNAs) were expressed as part of a mir30-based GIPZ lentiviral vector (Dharmacon). The clones used in this study include BAP1 (V2LHS_41473), DR4 (V3LHS_383718), DR5 (V3LHS_328891), YY1 (V3LHS_412955) and the empty GIPZ control vector.

### Lentivirus production and cell transfection

Lentiviral particles were produced by co-transfection of 293T cells with construct plasmids and the packaging plasmids pCMV-dR8.74 and pMD2.G (kind gifts from Dr Adrian Thrasher, UCL) using a DNA transfection reagent jetPEI (Source Bioscience UK Ltd). The viral particles were concentrated by ultracentrifugation at 17,000 rpm (SW28 rotor, Optima LE80K Ultracentrifuge, Beckman) for 2 hours at 4°C. To determine the titres of prepared lentivirus, 293T cells were transduced with serial dilutions of viruses in the presence of 8 µg/mL polybrene and protein expression was assessed by flow cytometry and immunoblotting.

### Flow cytometry

All flow cytometry analysis was performed on an LSR Fortessa analyser (Becton Dickinson). For analysis of BAP1 expression, cells were fixed, permiabiliszed and stained with primary antibody to BAP1 (c-4, Santa Cruz-SC28383; 1:50) and then with an AlexaFluor 488-conjugated anti-mouse antibody (Invitrogen A-21202; 1:200). For analysis of surface expression of DR4 and DR5, cells were stained with 1:100 dilution of PE-conjugated antibody (Biolegend; #307205 for DR4, #307405 for DR5, #400112 for isotype). FlowJo software was used to analyse all data.

### Quantitative RT-PCR

Total RNA was extracted from the cells using SV Total RNA Isolation System (Promega) according to the manufacture’s instructions. cDNA was synthesized using iScript Reverse Transcription Supermix for RT-qPCR (Bio-Rad). Quantitative PCR was performed using TaqMan probes (DR4: Hs00269492_m1, DR5: Hs00366278_m1, beta-2-microglobulin: Hs00187842_m1) and TaqMan Gene Expression Master Mix (Life Technologies Limited) as per the manufacture’s protocol. Relative expression of DR4 and DR5 was calculated using comparative CT method with a reference gene, beta-2-microglobulin.

### ChIP assay

The ChIP assay was carried out using EZ ChIP™ Chromatin Immunoprecipitation kit (Merck-Millipore) according to the manufacture’s instruction. Briefly, the cells were cross-linked, quenched and lysed then the chromatin was fragmented by sonication shearing. Protein/DNA complexes were diluted, pre-cleared with Protein G agarose beads, then immunoprecipitated (IP) by incubation with antibodies against BAP1 (Cell Signaling #78105), YY1 (Abcam #ab38422) or IgG (Cell Signaling #2729) overnight with rotation, followed by incubation with protein G agarose beads for 1 hr. After washing beads, protein/DNA complexes were eluted, reverse crosslinked to free DNA, which was then purified using spin columns and analysed by quantitative PCR (qPCR). Primer pairs for ChIP assays were as follows: DR5; forward 5’-GGGAAGGGGAGAAGATCAAG-3’, reverse 5’-GAAGGGACCGGAACTAACCT-3’. DR4; forward 5’-CCGAATGCGAAGTTCTGTCT-3’, reverse 5’-AAGAGCCCCACACTTTGCT-3’.

### Luciferase Reporter assay

Meso-8T cells were transduced with lentiviral vectors expressing a firefly luciferase reporter plasmid containing either DR4 promoter (upstream −1773/+63) or DR5 promoter (upstream −1400), plus control Renilla luciferase reporter under a control of CMV promoter (pDR4-FireflyLuc-CMV-RenillaLucDsRed2 or pDR5-FireflyLuc-CMV-RenillaLucDsRed2) vectors (44). Cells were seeded in 96 wells plate and luciferase activities were measured using Dual-Luciferase Reporter Assay System kit as described by the manufacture (Promega). Fluc/Rluc ratios were determined as relative luciferase activities.

### Immunohistochemistry (IHC)

Tumor biopsies taken from patients with MPM in the MS01 trial (NCT00075699) were stored as formalin fixed paraffin embedded (FFPE) blocks or as unstained mounted sections as previously described (37). The TMA slides containing tumour samples from patients with MPM were purchased from MesobanK UK. To assess expression of DR4, DR5, CK5 and calretinin, samples were first incubated in the oven at 60 °C for 30min, then deparaffinised and rehydrated using an automated tissue processor (Tissue-Tek, The Netherlands). Antigen retrieval was achieved by immersion in 10mM Citric acid buffer (pH.6.0) at 95 °C for 15 min. After washing with PBS and blocking with 2.5% normal goat serum, samples were incubated with primary antibody: anti-DR4 (abcam; ab8414, 1:500), anti-DR5, (abcam: ab8416, 1:500), anti-calretinin (Leica Biosystems: NCL-L-CALRET-566, 1:200), anti-keratin 5 (Biolegend: 905501, 1:500) in 1% BSA / 4% serum overnight at 4 °C. Samples were incubated with ImmPRESS polymer reagent (VECTOR Laboratories) for 30min and stained with ImmPACT Nova RED (VECTOR Laboratories). Hematoxylin and eosin (H&E) staining was carried out using an automated tissue processor (Tissue-Tek, The Netherlands). Staining for BAP1 was performed as described before using anti-BAP1 antibody (Santa Cruz Biotechnology, sc-28282, 1:150) (37). Images were acquired using a NanoZoomer 2.0HT whole slide imaging system (Hamamatsu Photonics, Japan). Histology and nuclear BAP1 assessment was performed by two consultant pathologists. Intensity of DR4 and DR5 expression was assessed blindly by three independent observers and scored as follows (no staining=0; low staining=1; medium staining=2; strong staining=3).

### Bioinformatical Analysis

To identify the common transcription factors which potentially regulate these genes, the 2000 nucleotide sequence of the promoter regions of DR4 and DR5 are entered into Human Core-Promoter Finder (http://rulai.cshl.org/tools/genefinder/CPROMOTER/human.htm).

### Statistical Analysis

Data were evaluated using the statistical analysis and indicated with *P* values. *P*<0.05 was considered statistically significant. Using Prism 8 (GraphPad, CA, USA), student’s *t*-test was performed to analyse differences between two groups whilst one-way ANOVA was used to determine the differences between three or more independent groups. For the statistical analysis of TMAs, linear mixed modelling was used to account for multiple samples per patient, including the patient ID as a random effect. Linear mixed models were implemented using the Bioconductor *Ime4* and *ImerTest* packages. Pairwise *t*-test confirmed that there was no systematic bias between the score of different observers.

## Acknowledgement

We would like to thank Dr. Rachel Chambers for providing human primary fibroblasts, Dr. Deepak Chandrasekaran and Dr. Pascal. Durrenberger for help with histological analysis (University College London Respiratory). This research was supported by Wellcome Trust (to S.M.J: WT107963AIA, to N.K:106555/Z/14/Z).

