## Supplemental data for "BAP1 and YY1 regulate expression of death receptors in malignant pleural mesothelioma"

### Supplementary Figure S1

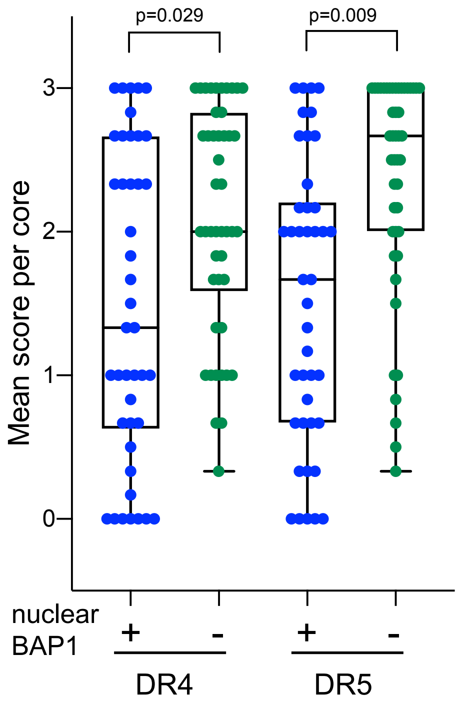


**Figure S1: DR4 and DR5 expression is inversely correlated with nuclear BAP1 expression in MPM TMA**

Semi-quantitative analysis of DR4 and DR5 expression in MPM TMA cores with (n=42) and without (n=46) nuclear BAP1 expression. Each dot represents an average score per core.  P-values are calculated using a linear mixed models, accounting for the patient ID as a random effect. BAP1 positive samples expressed less DR4 (p=0.029) and DR5 (p=0.0091)

### Supplementary Figure S2

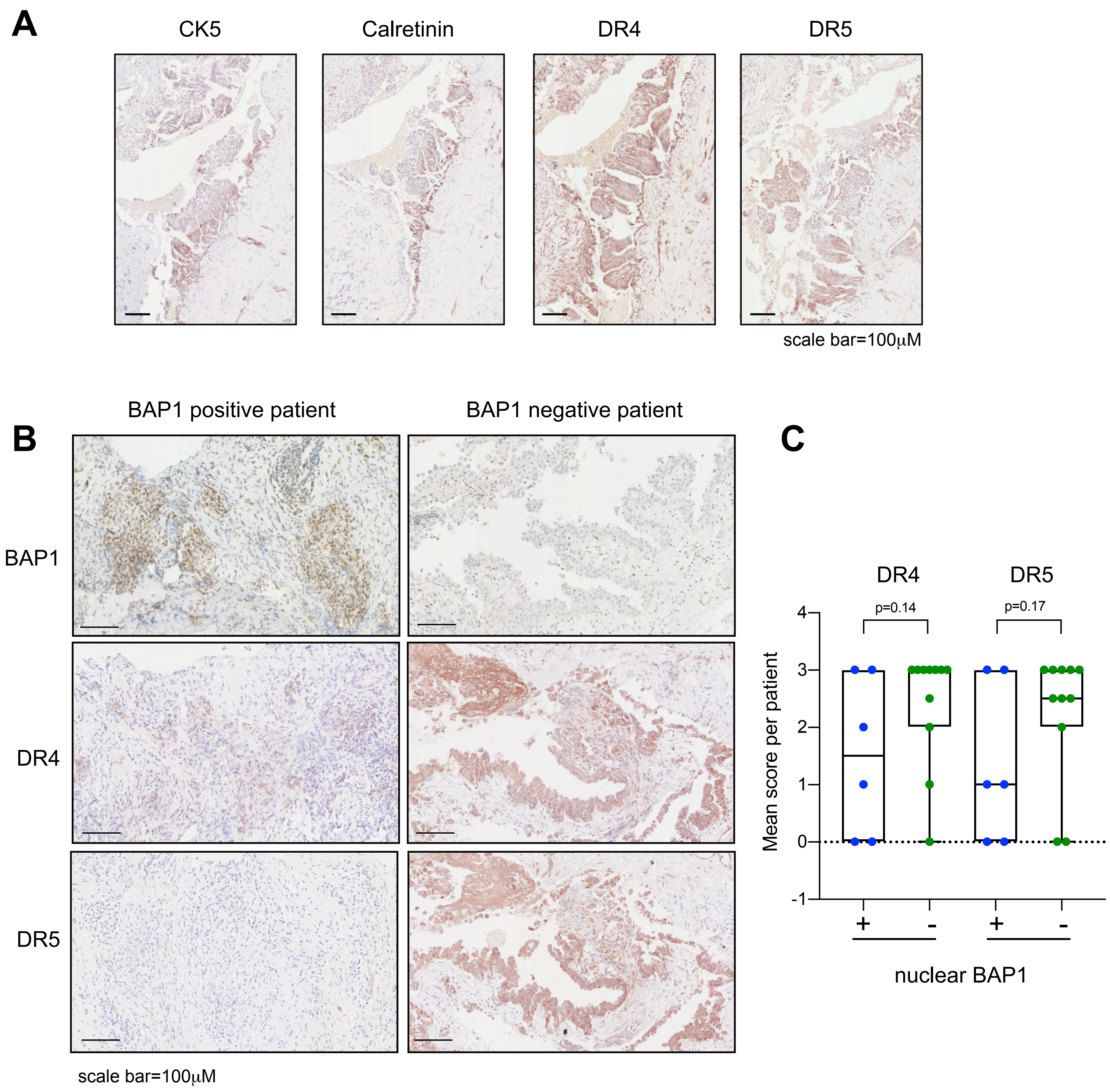


**Figure S2: Expression levels of DR4 and DR5 are inversely correlated with BAP1 expression in patients with MPM tumours**

**A**, Immunohistochemial analysis of DR4 and DR5 expression in primary MPM tissue. Two positive markers, cytokeratin 5 (CK5) and calretinin, were used to identify areas of MPM cells. Biopsy samples of 17 MPM patients (6 BAP1 positive, 11 BAP1-negative) from the MS01 clinical trial (NCT00075699) were used and representative images are shown. **B**, Immunohistochemical analysis of DR4, DR5 and BAP1 in primary MPM tissue as described in (A). Representative images show a strong inverse correlation between nuclear BAP1 expression and DR4/DR5 expression. **C**, Semi-quantitative analysis of IHC staining of DR4 and DR5 expression in primary MPM tissue with (+) (n=11) and without (-) (n=6) nuclear BAP1 staining. See method section for a detailed analysis.

### Supplementary Figure S3

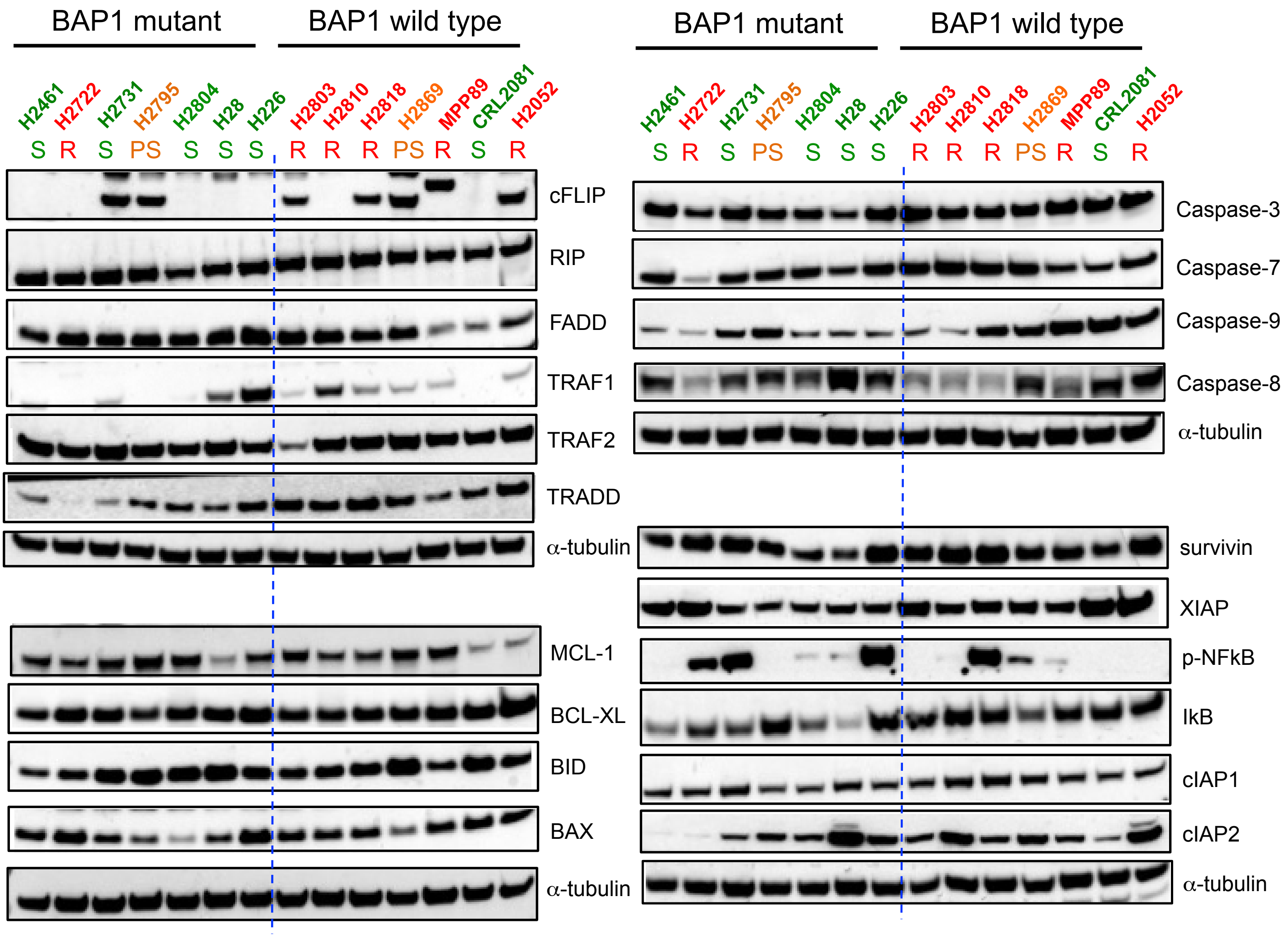


**Figure S3: Expression of proteins involved in TRAIL-induced signalling pathways in a panel of MPM cell lines**

MPM cell lines were divided into two groups (BAP1 mutant and BAP1 wild type) and are coloured according to the sensitivity to rTRAIL (green=sensitive (S); orange=partially sensitive (PS); red=resistant (R)). Proteins involved in TRAIL-induced signalling pathways, including pro- and anti-apoptotic pathways, were determined by immunoblots.

### Supplementary Figure S4

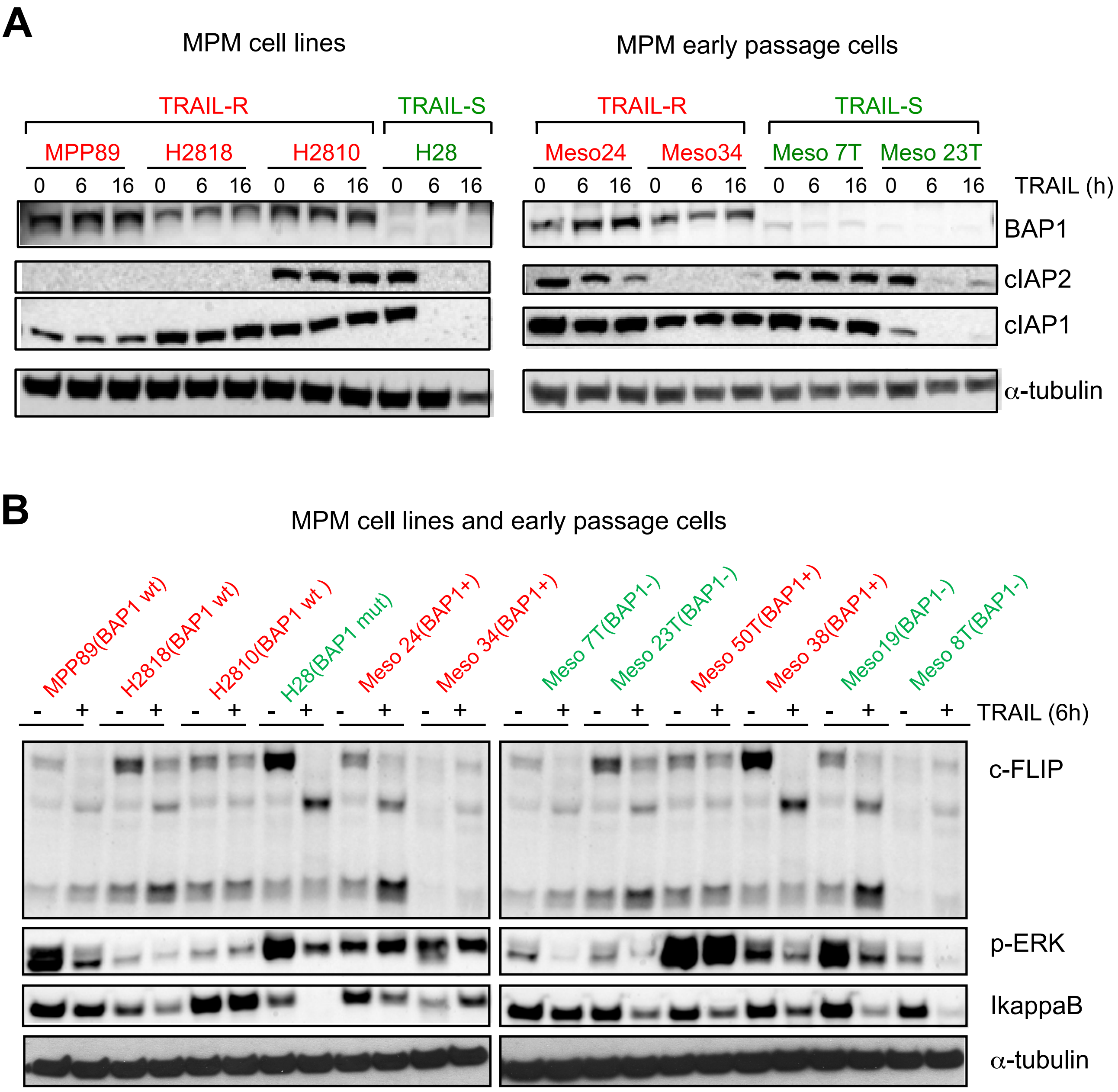


**Figure S4: rTRAIL-resistance in BAP1-expressing cells is not due to the induction of anti-apoptotic proteins by rTRAIL treatment.**

MPM cell lines and human early passage MPM cells were treated with 100ng/ml of rTRAIL for 0, 6 or 16 hours (**A**) or 0, 6 hours (**B**) to determine the expression of inhibitors of apoptosis proteins (cIAP1 and cIAP2), c-FLIP, p-ERK and IkappaB by western blot. Sensitivity to rTRAIL treatment is indicated as font colour: green sensitive (S); red resistant (R). BAP1's mutational status (wild type; wt or mutant; mut) or nuclear BAP1 expression (BAP1+ or BAP1-), which reflects the mutational status, is indicated (**B**).

### Supplementary Figure S5

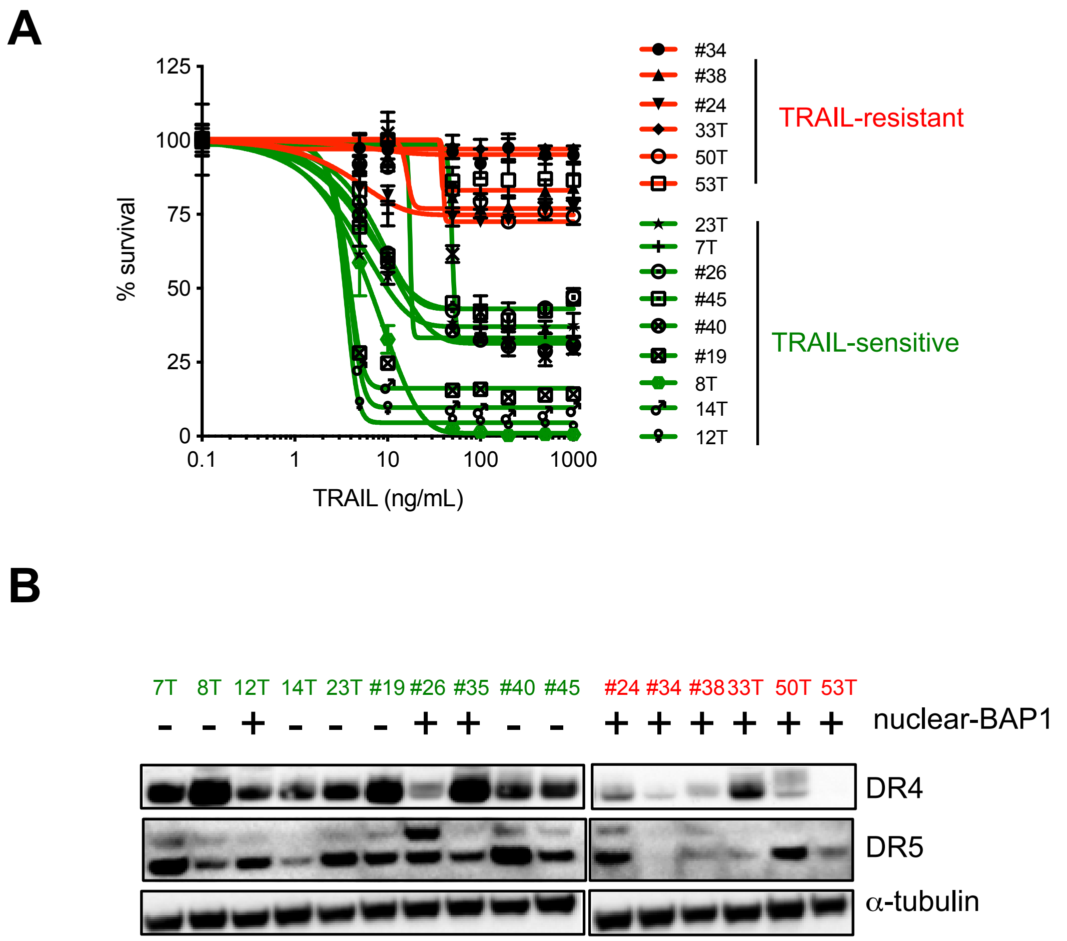


**Figure S5: Loss of nuclear BAP1 expression correlates with increased DR4 and DR5 expression and increased rTRAIL sensitivity in early passage MPM cells**

**A**, Cell viability assay of early passage MPM cells following treatment with a dose range of rTRAIL (0-1000ng/ml) for 72 hours. Cells with IC50>100ng/mL are defined as TRAIL-resistant (red); cells with IC50<100ng/mL are defined as TRAIL-sensitive (green). **B**, Immunoblots of DR4 and DR5 expression in early passage MPM cells stratified by rTRAIL sensitivity and the presence (+) or absence (-) of nuclear BAP1 expression Quantitative analysis of this experiment can be found in Fig.1E.

#### Supplementary Figure S6

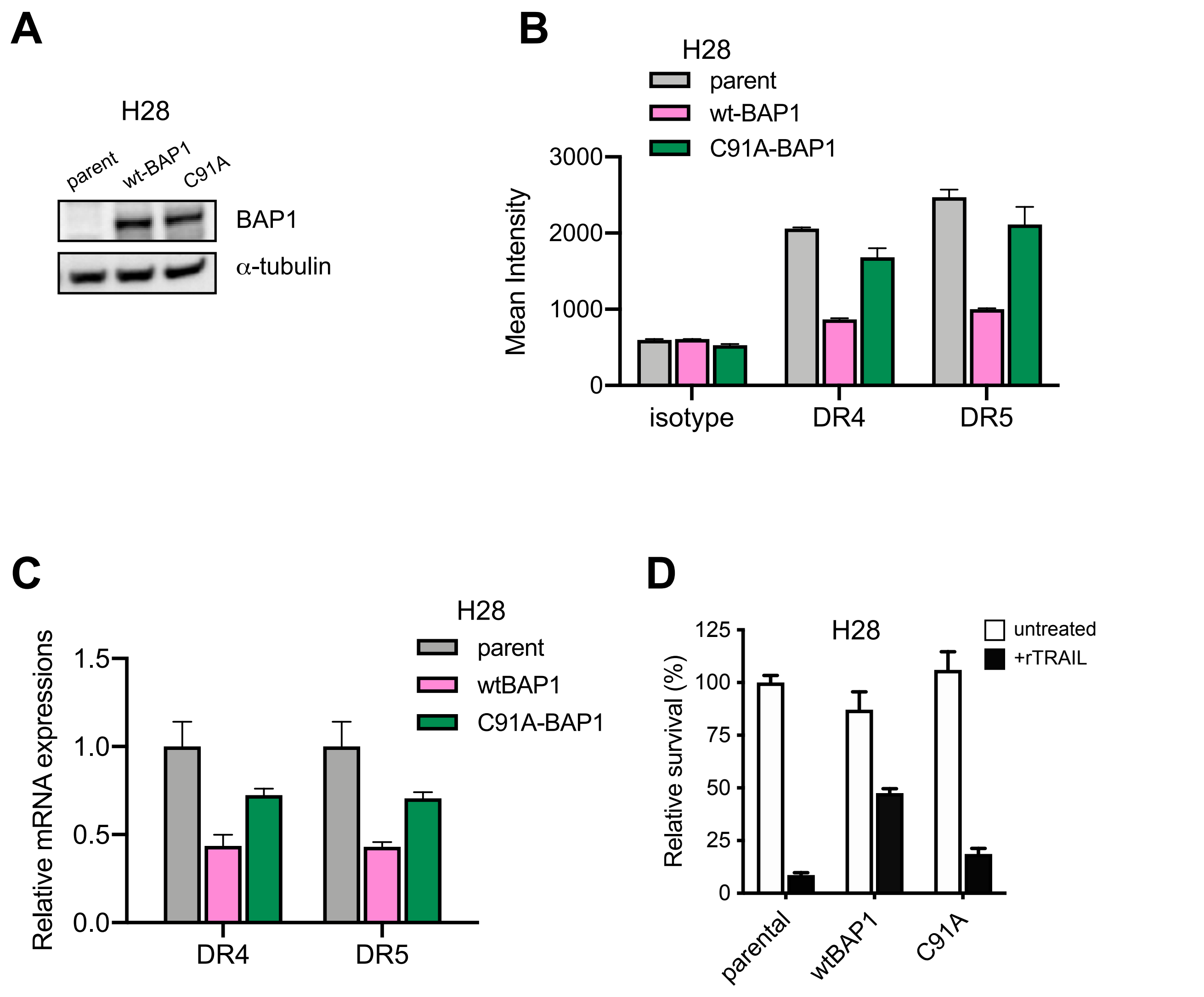


**Figure S6: BAP1 negatively regulates expression of DR4 and DR5 leading to TRAIL resistance in H28 MPM cells**

**A**, Immunoblot of BAP1 null H28 cells transduced with wild-type-BAP1 (wt-BAP1) or catalytically inactive BAP1-mutant (C91A-BAP1). **B**, Flow cytometry analysis of DR4 and DR5 cell surface expression in H28 cells transduced with wt-BAP1 or C91A-BAP1. Data shown are the mean ± s.d. performed in triplicates. **C**, Quantitative PCR analysis of DR4 and DR5 mRNA in H28 cells transduced with wt-BAP1 or C91A-BAP1. Data shown are the mean ± s.d. performed in triplicates. **D**, The sensitivity to rTRAIL (50ng/mL) is decreased in wt-BAP1-transduced H28 cells. Cell viability was assessed 72 hours later by XTT assay. Data shown are the mean ± s.d. (n=6).

### Supplementary Figure S7

| only DR4 | only DR5 | Both DR4&DR5 |
|---|---|---|
| NR3C1 | USF2 | YY1 |
| HNF1A | NF1 | NFKB1 |
| IRF2 | ETS2.1 | TP53 |
| USF2 | NFKB1.1 | FOXA1 |
| RARA | | CEBPB |
| ELF1 | | NFIC |
| RXRA.2 | | TAF6 |
| ARNT | | XBP1 |
| | | EBF2 |
| | | RXRA |
| | | RXRA.1 |
| | | NFYA |
| | | E2F1 |
| | | ATF3 |
| | | ETS2 |
| | | ETFB |
| | | STAT1 |
| | | POU2F1 |
| | | SP1 |
| | | JUN |
| | | GATA2 |
| | | LEF1 |
| | | RELA |
| | | ELK1 |

**Figure S7: A list of transcription factors that potentially bind to DR4 and/or DR5 promoters**

The promoter sequences, that are 2000 bases upstream of the coding region, were put into an online tool, Human Core-Promoter Finder (http://rulai.cshl.org/tools/genefinder/CPROMOTER/human.htm), to find transcription factors predicted to bind DR4 and/or DR5.

### Supplementary Figure S8

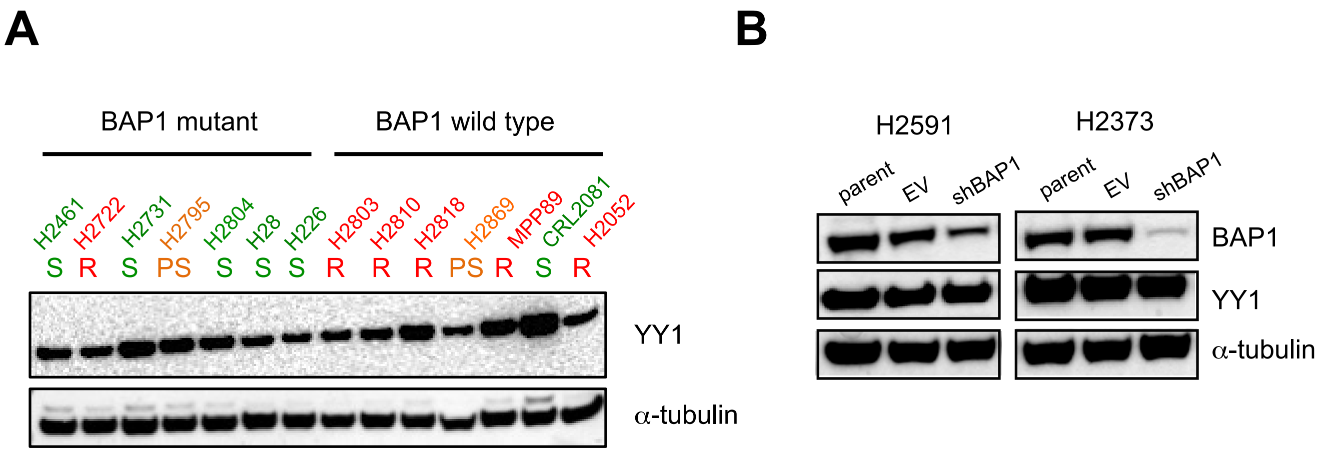


**Figure S8: YY1 expression is not regulated by BAP1**

**A**, Immunoblot of YY1 expression in BAP1 mutant (n=7) vs BAP1 wild-type (n=7) MPM cells. Sensitivity to rTRAIL treatment is indicated as font colour: green sensitive (S); orange partially sensitive (PS); red resistant (R). **B**, Immunoblot of YY1 expression in parental, BAP1 shRNA (shBAP1) or empty vector shRNA (EV) transduced H2591 and H2372 cells.

#### Supplementary Table1

#### List of antibodies used for immunoblotting

| primary antibody | secondary antibody | company & catalog number |
|---|---|---|
| BAP1 | Mouse | Santa Cruz Biotech #sc-28383 |
| BAX | Rabbit | Cell Signaling #5023 |
| Bcl-2 | Rabbit | Cell Signaling #2870 |
| Bcl-xL | Rabbit | Cell Signaling #2764 |
| BID | Rabbit | Cell Signaling #2002 |
| Bim | Rabbit | Cell Signaling #2819 |
| caspase 3 | Rabbit | Cell Signaling #9662 |
| caspase 7 | Mouse | Cell Signaling #9494 |
| caspase 8 | Mouse | Cell Signaling #9746 |
| caspase 9 | Rabbit | Cell Signaling #9502 |
| c-FLIP | Mouse | Enzo Life Science #ALX804-961 |
| c-IAP2 | Rabbit | Cell Signaling #3130 |
| c-IAP1 | Rabbit | Cell Signaling #7065 |
| PARP | Mouse | Cell Signaling #9546 |
| DR4 | Rabbit | Cell Signaling #42533 |
| DR5 | Rabbit | Cell Signaling #8074 |
| FADD | Rabbit | Cell Signaling #2782 |
| IkB (L35A5) | Mouse | Cell Signaling #4814 |
| Mcl-1 | Rabbit | Cell Signaling #5453 |
| phospho-NF-kN p65 (Ser536) | Rabbit | Cell Signaling #3033 |
| phospho-p42/44 MAPK(Erk1/2) | Rabbit | Cell Signaling #9101 |
| RIP | Rabbit | Cell Signaling #3493 |
| survivin | Rabbit | Cell Signaling #2803 |
| TRADD | Rabbit | Cell Signaling #3684 |
| TRAF1 | Rabbit | Cell Signaling #4715 |
| TRAF2 | Rabbit | Cell Signaling #4724 |
| XIAP | Rabbit | Cell Signaling #2045 |
| YY1 | Rabbit | Cell Signaling #2185 |
| alpha-tuublin | Rabbit, HRP-conjugated | Cell Signaling #9099 |

| secondary antibody | company & catalog number |
|---|---|
| anti-rabbit IgG, HRP-conjugated | Cell Signaling #7074 |
| anti-mouse IgG, HRP-conjugated | Cell Signaling #7076 |
